# Epigenetic regulation of a mouse PPARγ splicing variant, *Pparγ1sv*, during adipogenesis in 3T3-L1 cells

**DOI:** 10.1101/647842

**Authors:** Yasuhiro Takenaka, Takanari Nakano, Masaaki Ikeda, Yoshihiko Kakinuma, Ikuo Inoue

## Abstract

We have previously reported the abundant and ubiquitous expression of a newly identified splicing variant of mouse peroxisome proliferator-activated receptor-γ (*Pparγ*), namely *Pparγ1sv* that encodes PPARγ1 protein, and plays an important role in adipogenesis. *Pparγ1sv* has a unique 5’UTR sequence, compared to those of mouse *Pparγ1* and *Pparγ2* mRNAs. This implies the presence of a novel transcriptional initiation site and promoter for *Pparγ1sv*. We found that DNA methylation of 42 CpG sites in the proximal promoter region (−733 to −76) of *Pparγ1sv* was largely unchanged five days after adipocyte differentiation, whereas chromatin immunoprecipitation-quantitative PCR (ChIP-qPCR) using antibodies against H3K4me3 and H3K27ac revealed that these modifications significantly elevated at the transcription start sites of *Pparγ1sv* and *Pparγ2* after differentiation.

## Introduction

The peroxisome proliferator-activated receptors (PPARs) function as nuclear receptors to regulate the expression of many genes involved in metabolic homeostasis. PPARγ is the third member of PPAR subtype of genes and is one of the master regulators of adipogenesis^1^. In a ligand-dependent manner, PPARγ regulates transcription of target genes such as *Fabp4* and 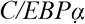which are indispensable for completion of adipocyte differentiation. There are two isoforms of PPARγ; the ubiquitously expressed PPARγ1 and the adipocyte-specific PPARγ2. PPARγ2 in mice is longer than PPARγ1 by 30 amino acid residues at the N-terminus. In to *Pparγ1* and *Pparγ2* mRNAs, we have recently reported a novel mouse *Pparγ* splicing variant, *Pparγ1sv*, that is indispensable for adipocyte differentiation of 3T3-L1 and mouse primary cultured preadipocytes^2^. *Pparγ1sv* was significantly up-regulated during adipocyte differentiation of 3T3-L1 cells and mouse primary cultured preadipocytes, and its inhibition by specific siRNAs completely abolished the induced adipogenesis in 3T3-L1 cells. *Pparγ1sv* has a unique 5’UTR sequence, implying the presence of a unique transcriptional initiation site and regulatory elements for the expression. Both the *Pparγ1sv* and *Pparγ2* promoters are transactivated by the overexpression of C/EBPβ and C/EBPδ, which are transiently expressed very early during adipocyte differentiation^3^, in 3T3-L1 cells. However, detailed mechanism of *Pparγ1sv* regulation during adipogenesis remains to be elucidated.

Methylated CpG sites on the *Pparγ2* promoter are progressively demethylated upon the induction of differentiation in 3T3-L1 cells^4^. In humans, a specific region of the *Pparγ1* promoter is methylated in colorectal cancers, which correlates with a lack of PPARγ expression^5^. In addition to DNA methylation of *Pparγ* promoters, histone modifications are involved in transcriptional regulation of *Pparγ* gene^6^. In this study, we report that epigenetic regulation of *Pparγ1sv* is via histone modifications, but not DNA methylation.

## Experimental procedures

### Cell culture and differentiation

3T3-L1 cells were obtained from JCRB Cell Bank (Osaka, Japan). Culture condition, media and method of adipocyte differentiation were described elsewhere^2^.

### Bisulfite sequencing

Genomic DNA was isolated from 3T3-L1 preadipocytes (day 0) and differentiated adipocytes (day 5) using DNeasy Tissue kit (Qiagen). Two µg of each DNA sample was bisulfite modified using EpiTect Bisulfite kit (Qiagen). The *Pparγ1sv* promoter region was amplified with EpiTaq HS polymerase (Takara) using the following primers: Novelpro_UP3bis, 5′-TGTGATAGATAAGGTGATAGAGTTTGG-3′ and Novelpro_LP1bis, 5′-TCCCTTATATAAAAACAACCCAAACTA-3′. PCR fragments were cloned into the pGEM-T Easy vector (Promega) and sequenced for both strands. Bisulfite sequences from all clones were analyzed using QUMA^7^.

### ChIP-qPCR analyses

3T3-L1 cells cultured in 10 cm dishes were fixed with 1% formaldehyde at around 25°C for 10 min, after which fixation was halted by addition of glycine solution. Immunoprecipitated protein/DNA complexes were prepared using the Magna ChIP A kit (Millipore) following manufacture’s instructions. Anti-acetyl-histone H3 (Lys27) and anti-monomethyl-histone H4 (Lys20) antibodies were purchased from Millipore. Anti-trimethyl-histone H3 (Lys4) antibody was purchased from Wako Pure Chemical Industries. ChIP samples were analyzed by qPCR with gene-specific primers as follows: NVpro_ChIP_UP1, 5′-GTCGGAGGGTGGGGAGGAGGATG-3′ and NVpro_ChIP_LP1, 5′-CCCAATCCCAAGCCATAAAGCAC-3′ for *Pparγ1sv* promoter; NVpro_ChIP_UP3, 5′-GGAGCAAGGCGGCCAGGTAACCA-3′ and NVpro_ChIP_LP3, 5′-GGCGGGTGCTGTGCGTCGGTGAG-3′ for *Pparγ1* promoter; g2pro_ChIP_UP2, 5′-GCCTTTATTCTGTCAACTATTCCTTTT-3′ and g2pro_ChIP_LP2, 5′-AGTATTTATCTTTGGTTGAAACTCCTA-3′ for *Pparγ2* promoter.

## Results

*Pparγ1sv* was remarkably up-regulated as early as day 3 upon induction of adipogenesis in 3T3-L1 cells^2^. To address how transcription of *Pparγ1sv* is regulated, we investigated the DNA methylation of CpG sites in the 658 bp upstream region of the *Pparγ1sv* transcription initiation site. The methylation status of 42 CpG sites in the −733/−76 of the *Pparγ1sv* promoter region (Fig. 1A, wave line) were analyzed by bisulfite genomic sequencing. Comparison of the methylation percentage before (day 0) and after (day 5) differentiation of 3T3-L1 cells revealed that 2 CpG sites at position −643 and −638 (Fig. 1B, two arrowheads) of differentiated 3T3-L1 were more frequently methylated than those of undifferentiated cells (P<0.05). However, the other 40 CpG sites were not significantly methylated or demethylated during adipocyte differentiation.

**FIGURE 1.**
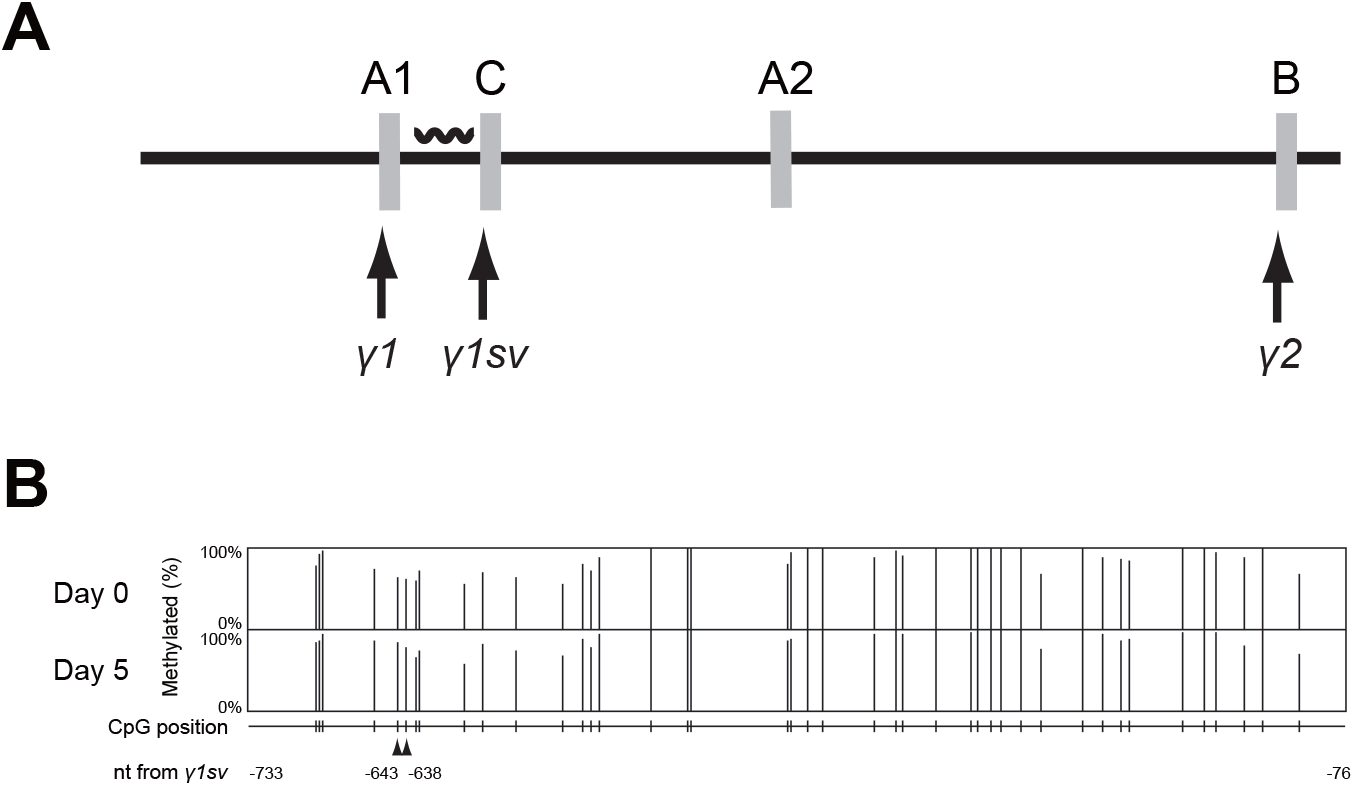
DNA methylation of *Pparγ1sv* promoter in adipogenesis. (A) Wave line denotes the region analyzed by bisulfite sequencing. Arrows indicate positions of transcription start sites of three *Pparγ* transcripts. (B) Percentages of DNA methylation of 42 CpG islands in the *Pparγ1sv* promoter (658-bp) were calculated by bisulfite sequencing of 90 (day 0) and 88 (day 5) clones. Arrowheads indicate CpG sites that were significantly methylated after the induction of adipocyte differentiation (day 5) as compared to undifferentiated cells (P<0.05).

Increase in levels of histone modification at the *Pparγ* promoter have been associated with adipocyte differentiation^8, 9^. We examined the trimethylation of histone H3 lysine 4 (H3K4me3), acetylation of histone H3 lysine 27 (H3K27ac), and monomethylation of histone H4 lysine 20 (H4K20me1) at the *Pparγ1sv* promoter (Fig. 2A). We found that H3K4me3 and H3K27ac are elevated in the region spanning the transcription start sites^10^. H4K20me1 increases in the downstream regions of transcription start sites for both *Pparγ1* and *Pparγ2*^8^. ChIP-qPCR using specific primers revealed remarkable increases in modifications by H3K4me3 and H3K27ac around the transcription start sites of *Pparγ1sv* and *Pparγ2* at day 5 of adipocyte differentiation (Fig. 2B). In contrast, H4K20me1 levels at the promoter regions of *Pparγ1sv* and *Pparγ2* were unchanged.

**FIGURE 2.**
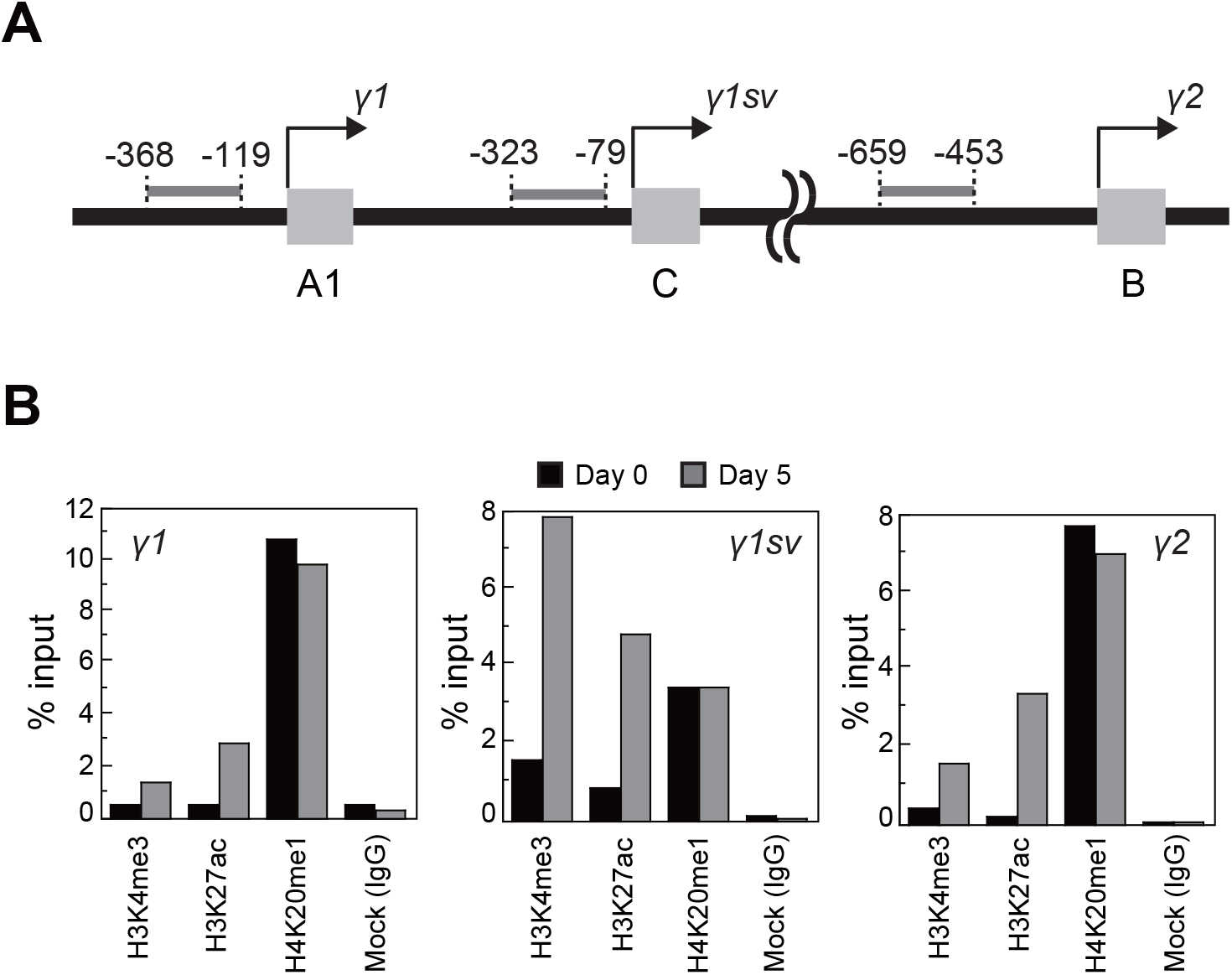
Histone modifications of *Pparγ1sv* promoter. H3K4me3, H3K27ac, and H4K20me1 were analyzed by ChIP-qPCR using specific primers for the promoter regions of *Pparγ1*, *Pparγ1sv*, and *Pparγ2*. (A) Arrows indicate positions of transcription start sites of the three transcription variants. Gray bar indicates analyzed promoter region. Distance from each transcription start site is indicated as numbers of nucleotides. (B) Enrichment of *Pparγ1sv* promoter region by pull-down with H3K4me3, H3K27ac, or H4K20me1 specific antibodies on day 5 of adipocyte differentiation. Values represent the mean of two experiments.

## Discussion

Previous studies have found that methylated CpG sites on the *Pparγ2* promoter get demethylated progressively upon adipogenesis in murine 3T3-L1^4^. Also, as mentioned before, studies on human colorectal cancer have shown an inverse correlation between methylation of specific regions of *Pparγ1* promoter and PPARγ expression^5^. In our studies, DNA methylation levels of the *Pparγ1sv* promoter were essentially unchanged during adipogenesis in 3T3-L1 cells (Fig. 1B). Therefore, epigenetic regulation by DNA methylation may not contribute to the regulation of *Pparγ1sv* expression. Instead, it is likely that *Pparγ1sv* and *Pparγ2* transcripts share a common regulatory mechanism by histone modifications. ChIP-qPCR using antibodies against H3K4me3 and H3K27ac revealed that these histone modifications significantly elevated at the transcription start sites of *Pparγ1sv* and *Pparγ2* by the induction of differentiation (Fig. 2B). H3K4me3 and H3K27ac levels on the *Pparγ1* promoter also increased but to a smaller extent relative to those of *Pparγ1sv* and *Pparγ2*, which may be indicative of a limited enhancement of *Pparγ1* mRNA expression during adipogenesis in 3T3-L1 cells.

## Acknowledgements

We are grateful to Ms. Sawako Sato (Saitama Medical University) for her technical assistances. We would like to thank Editage (www.editage.jp) for English language editing.

## Declaration of competing financial interests (CFI)

The authors declare they have no actual or potential competing financial interests.

## References

1. Tontonoz P, Spiegelman BM. Fat and beyond: the diverse biology of PPARγ. Annu. Rev. Biochem. 2008;77:289–312.

2. Takenaka Y, Inoue I, Nakano T, et al. A Novel Splicing Variant of Peroxisome Proliferator-Activated Receptor-γ (*Pparγ1sv*) Cooperatively Regulates Adipocyte Differentiation with *Pparγ2*. PLoS One. 2013;8:e65583.

3. Lefterova M, Lazar M. New developments in adipogenesis. Trends Endocrinol. Metab. 2009;20:107–114.

4. Fujiki K, Kano F, Shiota K, Murata M. Expression of the peroxisome proliferator activated receptor gamma gene is repressed by DNA methylation in visceral adipose tissue of mouse models of diabetes. BMC Biol. 2009;7:38.

5. Pancione M, Sabatino L, Fucci A, et al. Epigenetic silencing of peroxisome proliferator-activated receptor gamma is a biomarker for colorectal cancer progression and adverse patients’ outcome. PLoS One. 2010;5:e14229.

6. Lee JE, Ge K. Transcriptional and epigenetic regulation of PPARgamma expression during adipogenesis. Cell Biosci. 2014;4:29.

7. Kumaki Y, Oda M, Okano M. QUMA: quantification tool for methylation analysis. Nucleic Acids Res. 2008;36:W170–175.

8. Wakabayashi K, Okamura M, Tsutsumi S, et al. The peroxisome proliferator-activated receptor gamma/retinoid X receptor alpha heterodimer targets the histone modification enzyme PR-Set7/Setd8 gene and regulates adipogenesis through a positive feedback loop. Mol. Cell. Biol. 2009;29:3544–3555.

9. Waki H, Nakamura M, Yamauchi T, et al. Global mapping of cell type-specific open chromatin by FAIRE-seq reveals the regulatory role of the NFI family in adipocyte differentiation. PLoS Genet. 2011;7:e1002311.

10. Wang Z, Zang C, Rosenfeld JA, et al. Combinatorial patterns of histone acetylations and methylations in the human genome. Nat. Genet. 2008;40:897–903.

